# Concurrent lipidomics and proteomics on malignant plasma cells from multiple myeloma patients: Probing the lipid metabolome

**DOI:** 10.1101/702993

**Authors:** Ahmed Mohamed, Joel Collins, Hui Jiang, Jeffrey Molendijk, Thomas Stoll, Federico Torta, Markus R Wenk, Robert J Bird, Paula Marlton, Peter Mollee, Kate A Markey, Michelle M Hill

## Abstract

**Background:** Multiple myeloma (MM) is a hematological malignancy characterized by the clonal expansion of malignant plasma cells. Though durable remissions are possible, MM is considered incurable, with relapse occurring in almost all patients. There has been limited data reported on the lipid metabolism changes in plasma cells during MM progression. Here, we evaluated the feasibility of concurrent lipidomics and proteomics analyses from patient plasma cells, and report these data on a limited number of patient samples, demonstrating the feasibility of the method, and establishing hypotheses to be evaluated in the future.

**Methods:** Plasma cells were purified from fresh bone marrow aspirates using CD138 microbeads. Proteins and lipids were extracted using a bi-phasic solvent system with methanol, methyl tert-butyl ether, and water. Untargeted proteomics, untargeted and targeted lipidomics were performed on 7 patient samples using liquid chromatography-mass spectrometry. Two comparisons were conducted: high versus low risk; relapse versus newly diagnosed. Proteins and pathways enriched in the relapsed group was compared to a public transcriptomic dataset from Multiple Myeloma Research Consortium reference collection (n=222) at gene and pathways level.

**Results:** From one million purified plasma cells, we were able to extract material and complete untargeted (∼6000 and ∼3600 features in positive and negative mode respectively) and targeted lipidomics (313 lipids), as well as untargeted proteomics analysis (∼4100 reviewed proteins). Comparative analyses revealed limited differences between high and low risk groups (according to the standard clinical criteria), hence we focused on drawing comparisons between the relapsed and newly diagnosed patients. Untargeted and targeted lipidomics indicated significant down-regulation of phosphatidylcholines (PCs) in relapsed MM. Although there was limited overlap of the differential proteins/transcripts, 76 significantly enriched pathways in relapsed MM were common between proteomics and transcriptomics data. Further evaluation of transcriptomics data for lipid metabolism network revealed enriched correlation of PC, ceramide, cardiolipin, arachidonic acid and cholesterol metabolism pathways to be exclusively correlated among relapsed but not in newly-diagnosed patients.

**Conclusions:** This study establishes the feasibility and workflow to conduct integrated lipidomics and proteomics analyses on patient-derived plasma cells. Potential lipid metabolism changes associated with MM relapse warrant further investigation.

## Introduction

Multiple myeloma (MM) is an incurable plasma cell malignancy characterized by plasma cell infiltration of the bone marrow, and/or the presence of extramedullary plasmacytomas [2]. With an increasing number of treatment options available, median survival for MM has improved, and now approaches six years [5]. Despite advances in therapeutic strategies and an increasing number of pharmacological agents to choose from, MM eventually relapses for the majority of patients, hence there is a need to understand the mechanisms of relapse and identify potential new therapeutic approaches.

The Revised International Staging System (R-ISS) for MM incorporates serum biomarkers (lactate dehydrogenase, beta-2-microglobulin and albumin) and cytogenetic abnormalities of known prognostic significance to predict disease behavior [4]. It is imprecise however, with different patients in the same risk group exhibiting heterogeneous behavior and prognoses. MM treatment strategies predominantly use regimens built around immunomodulatory drugs such as thalidomide or its analogues, or proteasome inhibitors including bortezomib or carfilzomib. These treatments may be followed by autologous stem cell transplantation. With an increasing number of treatment options available, median survival has improved in the last decade, now approaching six years [5], but despite these advances, myeloma eventually relapses for the majority of patients.

Perturbations in lipid metabolism are emerging as potential drivers and therapeutic targets in cancer development and progression [7]. This is of particular relevance because obesity is a risk factor for a number of cancer types, including multiple myeloma (MM) [1]. A pooled analysis of 1.5 million participants from 20 unique prospective cohorts found a 1.2 to 1.5 fold increased risk of MM mortality with increasing body mass index [3]. In addition to the systemic chronic inflammation associated with obesity, increased bone marrow adiposity of the MM microenvironment may directly fuel MM progression [6].

In MM, initial lipidomic studies comparing malignant plasma cells to healthy plasma cells have reported decreased levels of phosphatidylcholines [8], and differing fatty acid composition of cellular membranes [8, 9]. There are limited studies on the metabolic changes that occur during MM relapse, with most studies focusing at the genomic level [10]. Using Raman spectroscopy to compare between drug resistant and sensitive MM cell lines, Franco *et al*. suggested differences in nuclear structure, as reflected by altered DNA:RNA ratio as well as cholesterol and phosphatidylethanolamine content [11]. Metabolic reprogramming, elevated oxidative stress response and up-regulated prostaglandin synthesis were reported by Zub *et al*. who compared the proteome and transcriptome of melphalan sensitive and resistant RPMI8226 cell lines [12].

Advances in omics technologies herald the potential of multi-omics systems analysis, where regulatory networks could be evaluated, for example, by combining proteomics and transcriptomics data. One challenge of performing multi-omics analysis on clinical samples is the limited patient-derived material. In this study, we investigated the feasibility of conducting lipidomic and proteomic analyses from the same patient-derived plasma cell sample. To validate the omics results from our pilot cohort, we compared the proteomics data with a larger public transcriptomic dataset from Multiple Myeloma Research Consortium reference collection, and interpreted the lipidomics data against a combined transcriptomics-proteomics lipid metabolism network for relapsed MM.

## Materials and Methods

### Study design and setting

A single-center, prospective pilot study was performed at the Princess Alexandra Hospital, Brisbane, Australia. We identified patients as possible candidates (on the basis of clinical features) prior to bone marrow aspiration and biopsy, and informed consent was sought prior to the aspiration and biopsy procedure. Bone marrow biopsies were all performed in the outpatient setting. Participant details are in Table S1.

### Plasma cell isolation from bone marrow

Plasma cells were isolated from fresh bone marrow aspirate samples using CD138 microbeads (Miltenyi). Purity was verified by flow cytometry (on the basis of CD38 and CD138 expression) and was >80% for all samples. Purified plasma cells were stored in aliquots of 10^6^ cells at −80°C until analysis.

### Lipid and protein extraction

Samples were selected based on laboratory confirmation of the diagnosis of myeloma with >10% plasma cells in the marrow aspirate sample, and >80% of CD138^+^ plasma cells post-purification. Extraction of lipids and proteins from 10^6^ isolated plasma cells was carried out using a bi-phasic solvent system of cold methanol, methyl tert-butyl ether (MTBE) and water [13]. Briefly, each sample was suspended in 20 µL of cold Milli-Q water and homogenized with a pipette tip, followed by addition of 20 µL of a 20 µM solution of zidovudine (AZT) in methanol as internal standard. Cold methanol (205 µL) was then added. The sample was vortexed briefly, frozen in liquid nitrogen for 2 min, thawed, and sonicated for 10 min. The freeze-thaw-sonication cycle was repeated twice. After incubating at −30 °C for 1 h, the sample was extracted by 750 µL cold MTBE with shaking at 4 °C for 15 min. Phase separation was induced by addition of 188 µL Milli-Q water, vortexing and centrifugation at 14000 g for 15 min at 4 °C. The upper phase was collected (700 µL) as the lipid-rich extract fraction, and protein was recovered as the pellet. The lipid extract was evaporated to dryness under vacuum and then reconstituted in 100 µL of a methanol/toluene (9:1) mixture for LC-MS analysis.

### Untargeted lipidomics

Untargeted lipidomics using LC-MS was performed as previously described [14], using Agilent 1290 Infinity II UHPLC with 6550 iFunnel Q-TOF mass spectrometer and Dual Agilent Jet Stream (AJS) source. Agilent Zorbax Eclipse Plus RRHD C18 column (2.1 × 50 mm, 1.8 µm) was used at a flow rate of 0.5 mL/min. Mobile phases for positive mode LC-MS consisted of A: acetonitrile/water (60:40) and B: isopropanol/acetonitrile (90:10). Both A and B contained 10 mM ammonium formate and 0.1% formic acid. In negative mode, ammonium formate and formic acid was replaced with 10 mM ammonium acetate in both eluents. LC gradient is described in **Supplementary File S2**.

Full scan MS spectra were acquired for samples at a mass range of m/z 100-1700. The TOF component was tuned using reference masses 118.09, 322.05, 622.03, 922.00, 1221.99 and 1521.97 in positive ionization mode, and the masses 112.99, 302.00, 601.98, 1033.99, 1333.97 and 1633.95 in negative mode. Source capillary voltages were set to 4000 V for positive ionization mode and 3500 V for negative ionization mode whilst the nozzle voltage was set to 0 V, fragmentor was set to 365 and octopoleRFPeak to 750. Nitrogen gas temperature was set to 250°C at a flow of 15 L/min and a sheath gas temperature of 400°C at a flow of 12 L/min. During the experiment reference masses were enabled for positive (121.05 and 922.01 Da) and negative modes (68.99, 112.98 and 1033.99 Da) to enable auto-recalibration of compounds with known masses.

The MS1 untargeted LC-MS data were subjected to batch Molecular Feature Extraction (MFE) with Agilent Profinder (B.08.00, Agilent Technologies Inc., Santa Clara, CA, USA). Data were then imported into R statistical framework for analysis. Data were first filtered to retained only features that in at least 75% of samples of one or more comparison groups. Remaining missing values were imputed with the minimum value. After quantile normalization and log2 transformation, differential analysis was carried out using limma package [15] to identify significant features (p value < 0.05, logFC > 1.5).

To assign the molecular identity to candidate features, LC-MS/MS was performed using nitrogen as the CID collision gas. MS/MS acquisition was performed in targeted mode. The HPLC, column and source parameters were identical to those used in the MS acquisition. A fixed collision energy of 20 eV was used to induce fragmentation for all targets in positive and negative mode. MS/MS data was acquired between 70-1500 m/z with MS and MS/MS scan rates of 3 spectra per second, with a maximum of 5 seconds between MS scans. The isolation width for all targets was set to medium (∼4 amu) and a delta retention time of 0.3 minutes. The LC-MS/MS data were submitted to the open source software MS-DIAL [16] with LipidBlast in-silico LC-MS/MS library [17] for identification of lipids.

### Targeted lipidomics

Targeted lipidomics experiments were performed using an Agilent Technologies 1290 Infinity II UHPLC system with an Agilent HILIC Plus RRHD 2.1×100 mm 1.8-micron column, coupled online to an Agilent 6490A Triple Quadrupole Mass spectrometer with iFunnel and AJS source. The mass spectrometer was operated in dynamic MRM mode. Each sample was analyzed in three separate dynamic MRM runs for the following lipid classes: Cer, PC and SM in method F1; PC-O, PC-P, HexCer, LPE, LPC in method F2; PE, PE-O, PE-P, PI, PG in method A1. MRM lipid transitions are shown in **Supplementary File S2**.

The source nitrogen gas temperature was set to 250°C at a flow of 15 L/min. The sheath gas temperature was 400°C with a flow of 12 L/min. The capillary voltage was set to 4000 V for positive mode and 5000 V for negative mode and the nebulizer operated at 30 psi. Ion funnel low and high pressure in positive mode were 150 and 60, and in negative mode 150 and 120 respectively. The autosampler was operated at 4°C and the column compartment was operated at 30°C for the duration of the experiment. A solution of 95% acetonitrile was used to perform the needle wash with a duration of 15 seconds. An injection volume of 8µL was used for all samples. Pooled quality control (QC) samples were injected multiple times to condition the HPLC column prior to analyzing the biological samples. Chromatographic separation of lipids was performed using 2 different HILIC buffer systems; 25 mM ammonium formate (pH4.6) or 10 mM ammonium acetate (pH7.6). The acetonitrile gradient was from 50% to 95% as described in **Supplementary File S2**.

Raw LC-MS data was imported into Skyline [18], where peak integration was automated but manually confirmed and adjusted if required. Retention time for internal standard of each lipid class was used to confirm correct peak integration of lipids belonging to the same class. Peak areas were exported from Skyline for further analysis in R. Data were then normalized using probabilistic quotient normalization [19] to correct for injection variations, and then log2 transformed. Differential analysis was carried out using limma package identify significant lipids (p value < 0.05, logFC > 1.5).

To perform enrichment analysis, lipid sets were generated based on class, total chain length and total chain unsaturation. Lipid set enrichment analysis was performed in R using the fgsea package [20].

### Proteomics

Proteins pellets were thawed on ice then centrifuged. Any excess liquid was removed and samples dried under N_2_ for 10 min. Protein pellets were resuspended in 15 uL of buffer (70 mM Tris pH8, 1% sodium deoxycholate, 10 mM tris(2-carboxyethyl)phosphine and 40 mM 2-chloroacetamide), and sonicated in the Bioruptor (Diagenode) for 15 minutes. Protein concentration was measured using DirectDetect® infrared spectrometer (Merck). A 10 μg aliquot of 1 mg/mL protein extract was denatured by heating at 95°C for 5 minutes. After cooling to room temperature, 0.2 μg trypsin (Promega) was added and incubated at 37C overnight. Digest was stopped by acidification to 0.5 % TFA, and peptides were isolated using OMIX C18 tips (Agilent). NanoLC-MS/MS was performed using a Waters nanoACQUITY UPLC system interfaced to an LTQ-Orbitrap Elite hybrid mass spectrometer as described in [21].

Acquired data was searched using MaxQuant [22] version 1.5.8.3 against SwissProt human proteome downloaded on 25/10/2017, and later exported to R for analysis. Proteins were filtered according to unique peptides (≥2) and Score (>5), and then according to missing values, where proteins were only kept if they were detected in at least 75% of samples of one or more comparison groups. Data was then quantile normalized and remaining missing values imputed using two techniques: i) proteins missing in < 25% of all samples were considered missing at random, and were imputed using localized least square regression as described in [23], ii) proteins missing in > 25% were imputed from a normal distribution centred at minimum intensity. Log2 transformed data was analyzed using limma package to identify significant proteins (p value < 0.05, logFC > 1.5). Pathway enrichment analysis was carried out using the fgsea package and pathways from Reactome database [24].

### Transcriptomics data set

Gene expression profiles of the Multiple Myeloma Research Consortium (MMRC) reference collection were downloaded from the Multiple Myeloma Genomics Portal (http://portals.broadinstitute.org/mmgp/) as a GCT file. Expression signals were obtained as median centered and log2 transformed, and imported into R. Patient samples were filtered to include only those diagnosed with Multiple myeloma and reported treatment status. Microarray probes were first mapped to UniProt IDs, followed by differential analysis and pathway enrichment using limma and fgsea packages, respectively.

### Network analysis

Biopax level 3 file of the “Metabolism of Lipids” pathway was downloaded from the Reactome database, imported and analyzed in R using NetPathMiner package [25]. Transcriptomic data was used to weight network based on adjacent pairwise correlation. Top 50 correlated paths, with a minimum path length of 6 reactions, were then extracted for relapsed and newly-diagnosed patients. Association of extracted paths with disease status was assessed by a path classification model. A subnetwork of top paths was then exported to Cytoscape [26] for interactive visualization and analysis.

## Results

Following clinical diagnosis, plasma cell isolation and quality control, a total of 7 participant samples were available for inclusion (Table S1). For each participant, 1×10^6^ plasma cells were extracted for proteomics and lipidomics analyses. Lipidomics was performed using both untargeted and targeted approaches. Two comparisons were conducted based on clinical information, with the caveat that the sample sizes were small in this study. Firstly, high risk MM (n=3) were compared to low risk MM (n=4) according to R-ISS staging. Secondly, relapsed/refractory MM (RRMM, n=2) versus newly diagnosed MM (NDMM, n=7). Table 1 summarizes the number of detected, filtered, and significant features for each analysis.

**Table 1.**
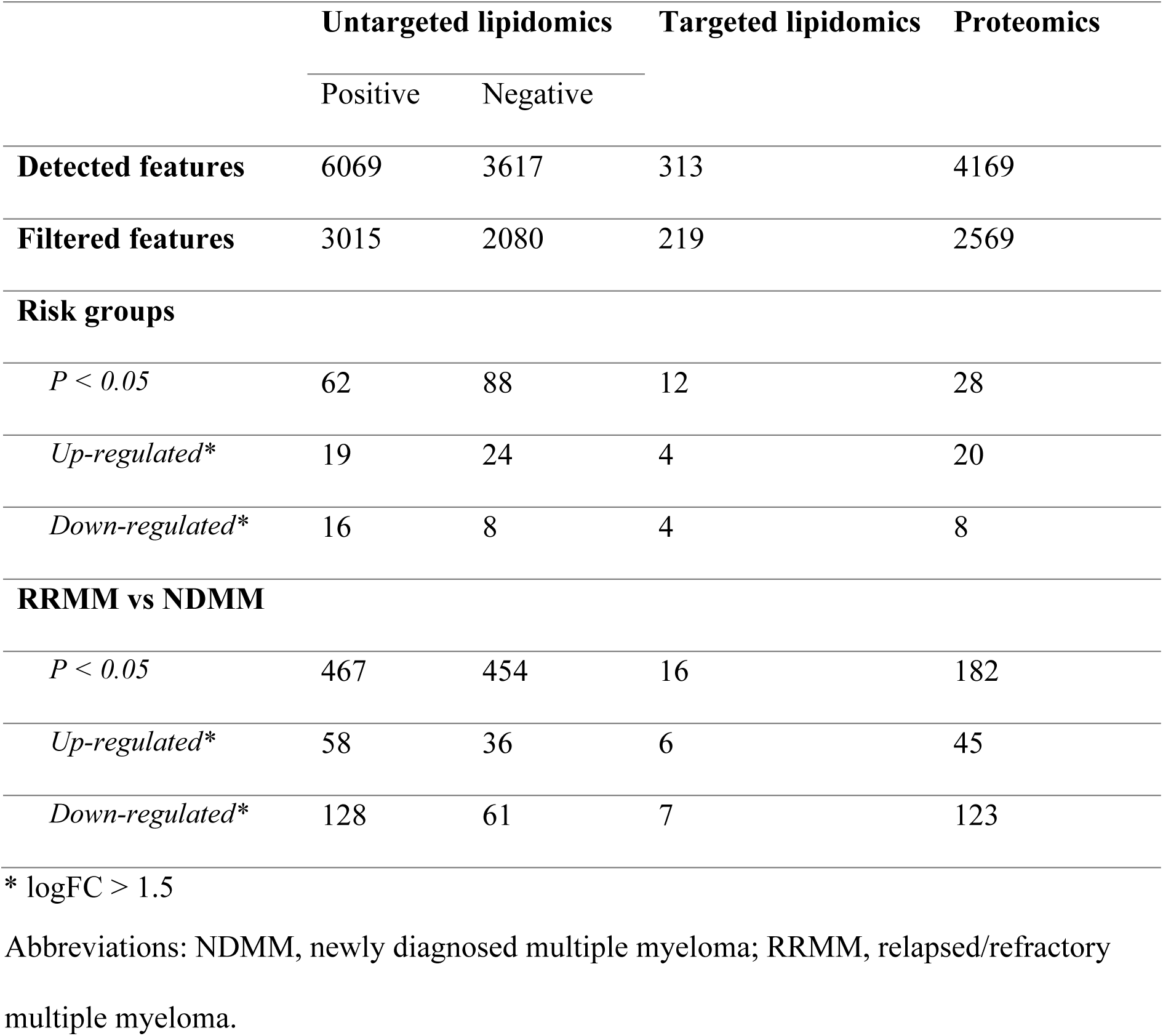
Overview of lipidomics and proteomics LC-MS analyses.

|  | Untargeted lipidomics |  | Targeted lipidomics | Proteomics |
| --- | --- | --- | --- | --- |
|  | Positive | Negative |  |  |
| **Detected features** | 6069 | 3617 | 313 | 4169 |
| **Filtered features** | 3015 | 2080 | 219 | 2569 |
| **Risk groups** |  |  |  |  |
| *P* < 0.05 | 62 | 88 | 12 | 28 |
| *Up-regulated\** | 19 | 24 | 4 | 20 |
| *Down-regulated\** | 16 | 8 | 4 | 8 |
| **RRMM vs NDMM** |  |  |  |  |
| *P* < 0.05 | 467 | 454 | 16 | 182 |
| *Up-regulated\** | 58 | 36 | 6 | 45 |
| *Down-regulated\** | 128 | 61 | 7 | 123 |

\* logFC > 1.5
Abbreviations: NDMM, newly diagnosed multiple myeloma; RRMM, relapsed/refractory multiple myeloma.

### Untargeted lipidomics profiling of plasma cells

For untargeted lipidomics profiling, 6069 and 3617 features were detected in the positive and negative mode, respectively. Filtering missing and low intensity features retained 3015 and 2080 features. Differential analysis between risk groups identified 62 and 88 significant features in positive and negative mode (**Supplementary File 3**). The number of significant features was much higher (>400 features) in RRMM/NDMM comparison, indicating higher variation compared to different risk groups. Differential features with logFC > 1.5 were selected for identification via MS/MS fragmentation and database matching using MS-DIAL. Out of ∼400 features, MS-DIAL matched 17 features to their lipid composition, in which several PCs were diminished in RRMM as well as in high risk patients (**Table 2**).

**Table 2.** Untargeted lipid features identified via MS/MS fragmentation.

| Lipid Molecule | ESI Mode | Comparison* | logFC |
| --- | --- | --- | --- |
| Cer[NS] 34:1; Cer[NS](d18:1/16:0); [M+H]<sup>+</sup> | + | high.low | 1.58 |
| Cer[NS] 34:2; Cer[NS](d18:1/16:1); [M+H]<sup>+</sup> | + | RRMM.NDMM | -2.90 |
| PC 30:0; [M+H]<sup>+</sup> | + | high.low | -1.75 |
| PC 30:0; [M+H]<sup>+</sup> | + | RRMM.NDMM | -1.66 |
| PC 31:1; [M+H]<sup>+</sup> | + | high.low | -1.87 |
| PC 31:1; [M+H]<sup>+</sup> | + | RRMM.NDMM | -1.56 |
| PC 32:2; [M+H]<sup>+</sup> | + | RRMM.NDMM | -1.56 |
| PC 34:4; [M+H]<sup>+</sup> | + | high.low | -2.39 |
| PC 34:4; [M+H]<sup>+</sup> | + | RRMM.NDMM | -2.10 |
| PC 35:4; [M+H]<sup>+</sup> | + | RRMM.NDMM | -1.67 |
| PC 40:4; [M+H]<sup>+</sup> | + | RRMM.NDMM | -3.82 |
| PC 40:7; [M+H]<sup>+</sup> | + | RRMM.NDMM | -1.55 |
| Plasmenyl-PC 30:0; [M+H]<sup>+</sup> | + | RRMM.NDMM | -3.83 |
| Plasmenyl-PC 36:1; [M+H]<sup>+</sup> | + | RRMM.NDMM | -2.55 |
| Plasmenyl-PC 38:5; [M+H]<sup>+</sup> | + | RRMM.NDMM | -4.12 |
| Plasmenyl-PE 40:6; [M-H]<sup>-</sup> | - | RRMM.NDMM | -1.76 |
| PS 36:4; [M+H]<sup>+</sup> | + | RRMM.NDMM | -2.24 |

\*Comparison between high and low risk group (high.low) or between relapse and newly-diagnosed (RRMM.NDMM)
Abbreviations: ESI, electrospray; logFC, log fold change

### Targeted lipidomics profiling of plasma cells

The targeted lipidomics method included 313 lipids, from which 219 lipids were retained after manual inspection and filtering through Skyline (**Supplementary File 4**). Differential analysis confirmed untargeted profiling results with several PCs diminished in both high risk and RRMM (**Table 3**). To investigate if the observed differences are specific to particular lipid class, we performed Lipid set enrichment analysis (**Figure 1, Supplementary File 4**), which revealed significant down-regulation trend in PCs in both high risk and RRMM. Ceramides and lyso-PEs were significantly enriched for upregulation in high risk patients, while down-regulated in RRMM. Elevated levels of phosphatidylethanolamines (PEs), sphingomyelins and sphingosines resulted in significant enrichment of these classes in RRMM.

**Table 3.** Reduced abundance of phosphatidylcholines (PC) in high risk and RRMM, measured by targeted lipidomics.

| Lipid Molecule | Comparison* | logFC |
| --- | --- | --- |
| PC 30:0 | high.low | -1.57819 |
| PC 30:1 | high.low | -1.37218 |
| PC 34:4 | high.low | -1.7996 |
| PC 34:4 | RRMM.NDMM | -1.14782 |
| PC 34:5 | high.low | -2.19233 |
| PC 38:0 | RRMM.NDMM | -3.03733 |
| PC 38:1 | RRMM.NDMM | -1.99746 |
| PC 40:0 | RRMM.NDMM | -2.39816 |
| PC 40:1 | RRMM.NDMM | -1.7426 |
| PC 40:2 | RRMM.NDMM | -1.13424 |
| PC(O-38:6) / PC(P-38:5) | RRMM.NDMM | -1.93524 |
| PC(O-40:7) / PC(P-40:6) | RRMM.NDMM | -1.77034 |

\*Comparison between high and low risk group (high.low) or between relapse and newly-diagnosed (RRMM.NDMM)
Abbreviations: logFC, log fold change

**Figure 1.**
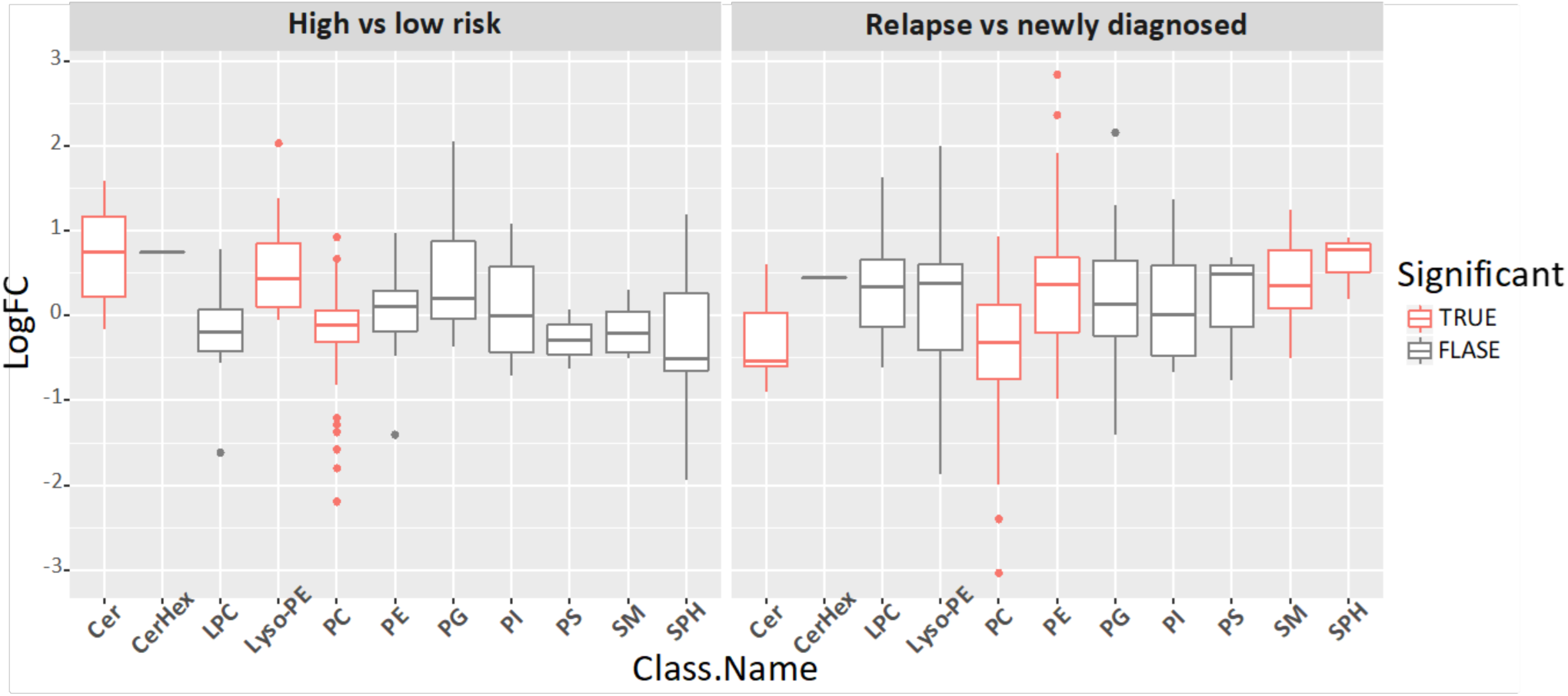
Targeted lipidomics measurements per lipid class, with significantly enriched classes marked with red. Targeted lipidomics data were grouped by lipid class and then evaluated for significance for high versus low risk MM (left) and RRMM versus NDMM (right) using enrichment analysis of fgsea R package. Lipid classes with adjusted P value < 0.05 are considered significantly different between the two groups (labelled red). LogFC, log fold change.

### Untargeted proteomics of plasma cells

In the untargeted proteomic analysis, 4169 proteins were identified, of which 2569 were subjected to differential analysis after filtering. Difference between risk groups was limited to 28 significant proteins, while RRMM vs NDMM comparison reported 182 differential proteins, the majority of which are down-regulated (**Supplementary File 5**). Enrichment analysis using Reactome pathways identified ∼ 150 significant pathways in RRMM (**Supplementary File 6**). In contrast, risk groups had only ∼25 enriched pathways, mostly related to extracellular matrix.

### Comparison of RRMM proteomics dataset with gene expression data

In both lipidomics and proteomics measurements, the differences between RRMM and NDMM were larger than those observed between risk groups. We followed up on these observations in RRMM by integrative analysis with the publicly available MMRC reference collection which contains gene expression profiles for plasma cells from a total of 222 patients, with 107 being NDMM (termed untreated) and 115 RRMM (termed treated). Mapping microarray probes to their corresponding UniProt IDs obtained expression levels for ∼ 17,000 genes. Differential expression analysis followed by pathway enrichment identified 430 significant pathways (**Supplementary File 6**).

There was significant overlap between the proteomics results from our cohort and the independent transcriptomics results at the pathway level but not at the gene level (hypergeometric test, **Figure 2**). Out of 6900 significantly expressed genes, 62 genes were also found significant at the protein level, only 20 of which were regulated in the same direction (p = 0.99) (**Supplementary File 5**). Interestingly, out of the 430 significantly enriched pathways in the transcriptomics dataset, 76 pathways were also enriched at the protein level, 67 of which in the same direction (p < 1e-16). Overlapped pathways included TCR, NF-kß signalling and protein synthesis pathways (**Supplementary File 6**).

**Figure 2.**
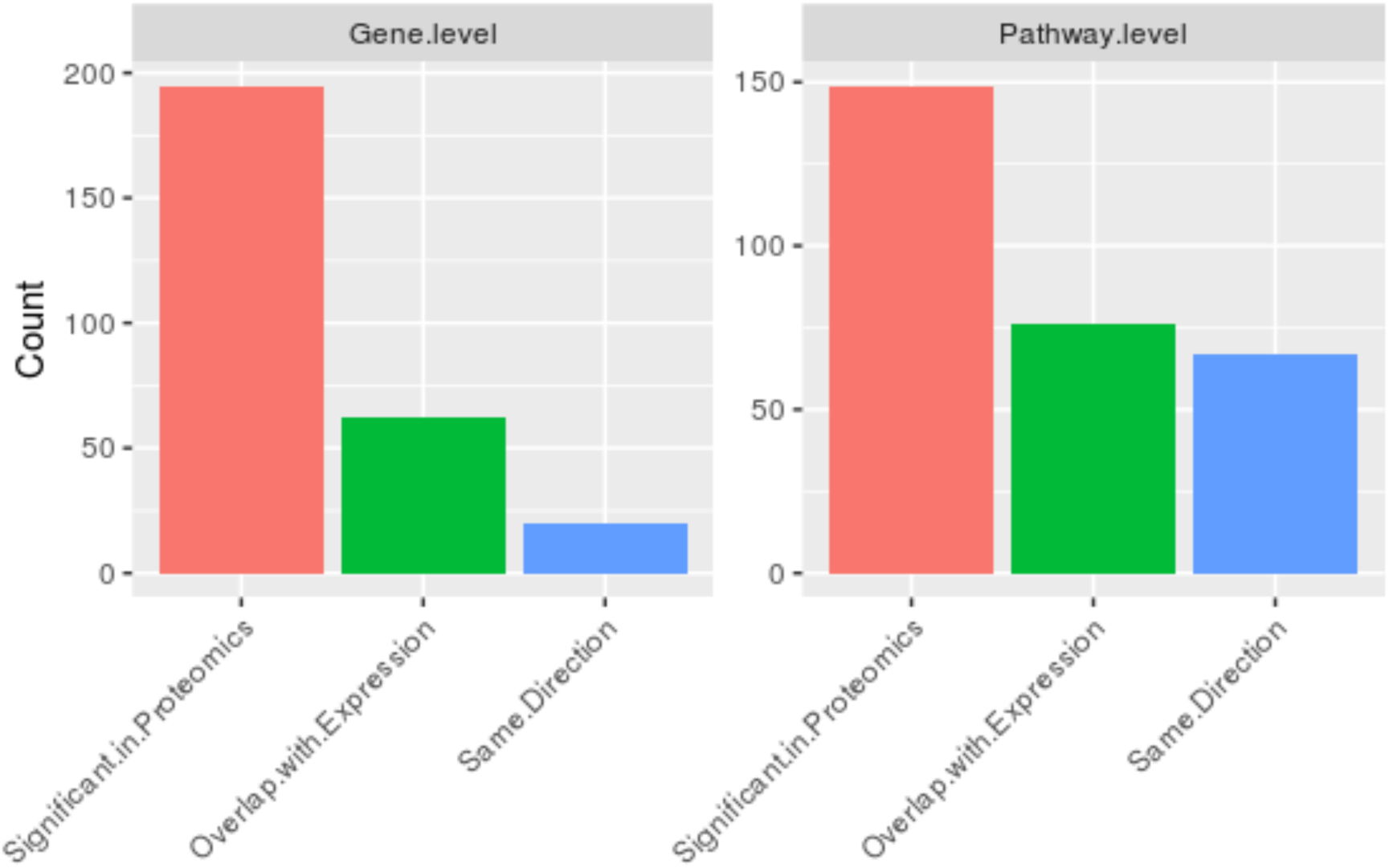
Overlap between proteomics and transcriptomics data at the gene and pathway levels. Proteomic level changes in RRMM compared to NDMM were evaluated against independent transcriptome data from the Multiple Myeloma Research Consortium reference collection. The graph shows the number of genes/proteins (left) or pathways (right) that are significantly different in the proteomics data (red bar), which also was significantly different in the transcriptome data (green bar), in the same direction (blue bar).

### Network analysis

Next, we focused on the lipid related pathways in RRMM. Reactome pathway group “Metabolism of lipids” was converted into a single connected network using NetPathMiner R package. Following the package instructions, small ubiquitous compounds, such as water and co-factors, were removed to prevent over-connectivity of the network, resulting in a network with 1130 nodes and 1571 edges. Metabolite nodes were then removed to obtain a reaction network, subsequently weighting the edges using transcriptomics datasets (see Methods). Top correlated paths showed strong association with their corresponding conditions. This was demonstrated by the ability of pathClassifier function to correctly predict path condition. Receiver Operating Characteristic (ROC) curve showed area under the curve (AUC) of 0.995, indicating high sensitivity and specificity of the path classifier (**Figure**).

Subnetworks constructed from correlated paths resulted in substantially smaller networks. In RRMM, a subnetwork of 101 nodes and 125 edges was obtained, with paths related to PCs, ceramides, cardiolipin metabolism, production of leukotrienes, exotoxins from arachidonic acid (AA), and production of dihydroxycholestanoic acid from cholesterol (**Figure 4**, red edges). On the other hand, the subnetwork correlated amongst NDMM consisted of 87 nodes and 96 edges, and incorporated FA and PE metabolism, production of prostaglandins and thromboxanes from AA, and production of phosphoserine from cholesterol. Subnetworks from both conditions showed a small overlap, with only 32 nodes and 23 edges (**Figure 4**, grey edges).

**Figure 3.**
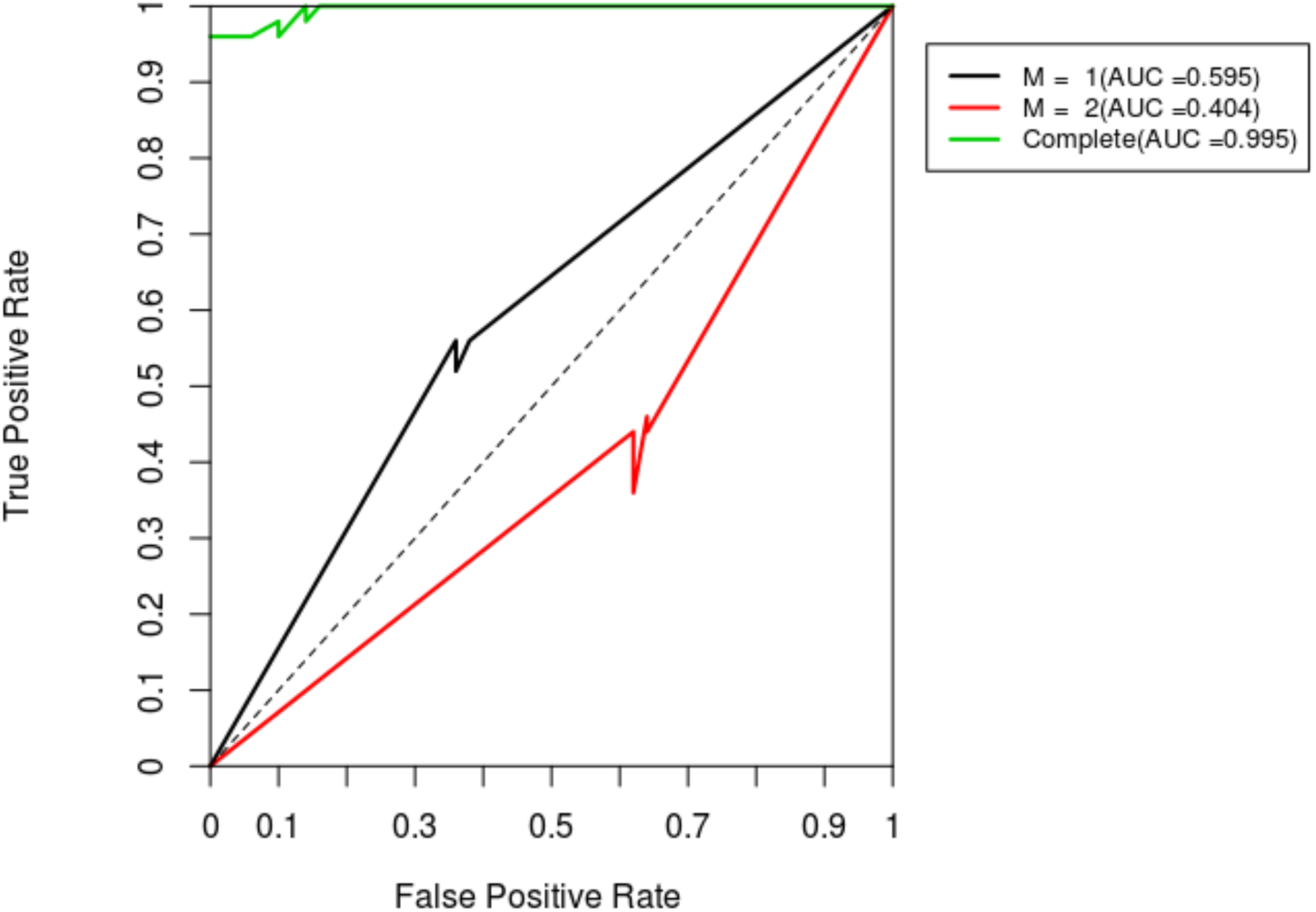
Receiver Operating Characteristic (ROC) curve for correlated path classification model of lipid metabolic pathways based on transcriptome data for RRMM. Diagnostic plot of the result from the path classification model for RRMM transcriptome data. ROC curves are shown for each component (M1, M2), which represent a path structure pattern. This gives information about which components is associated with RRMM and NDMM. A ROC curve with an AUC < 0.5 relates to RRMM. Conversely, ROC curve with AUC > 0.5 relates NDMM. Complete ROC represents the performance of the classifier using both components.

**Figure 4.**
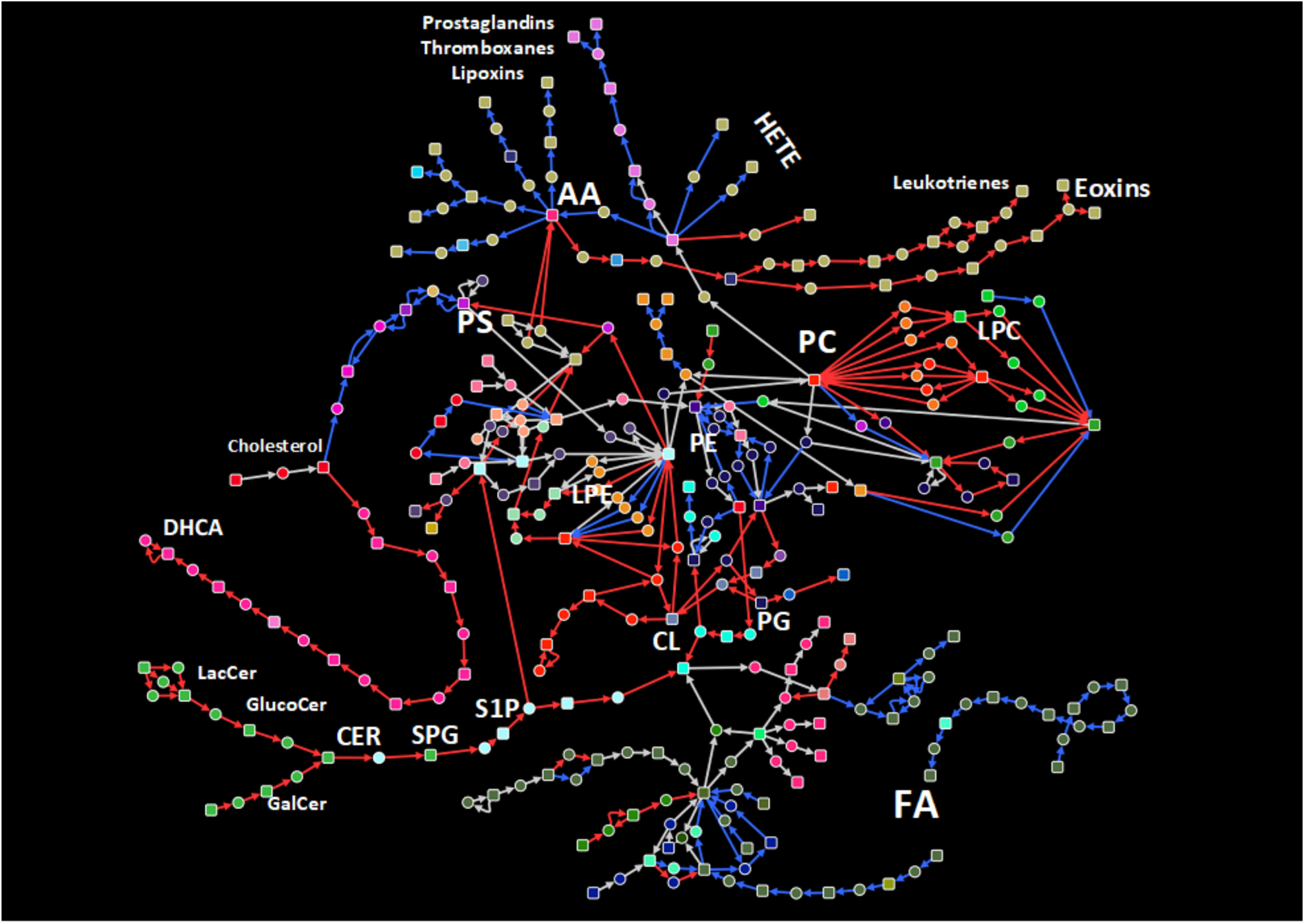
Extracted correlated lipid metabolism path network for RRMM and NDMM patients. A sub-network comprised of top 50 correlated paths based on gene expression in RRMM and NDMM was extracted from the lipid metabolism path network. Red and blue edges indicate exclusive correlation in RRMM and NDMM patients, respectively. Grey edges indicate correlation in both conditions.

Exploring the proteomics data in the context of correlated subnetwork for RRMM revealed a low detection rate (**Figure 5**). Notably, PLBD1, a phospholipase B implicated in sn1 and sn2 hydrolysis PCs, was up-regulated in RRMM proteomics and transcriptomics. This up-regulation of PLBD1, along with the correlation of PC metabolic subnetwork in RRMM, propose a possible explanation for the reduced levels of PCs observed in lipidomics data.

**Figure 5.**
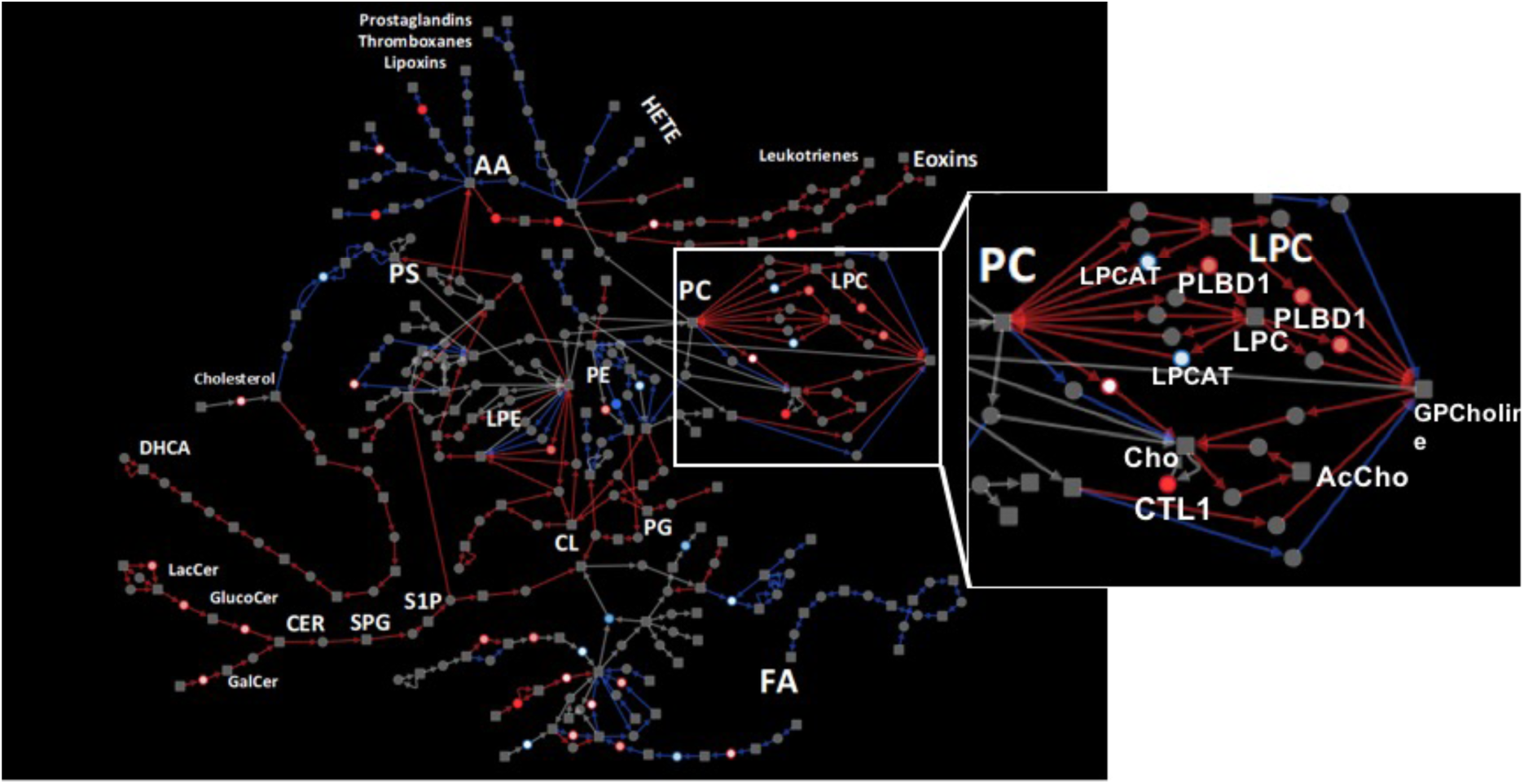
Proteomics results shown in the context of extracted lipid metabolism path network for RRMM and NDMM patients. Proteomic data were projected on to the same network shown in Figure 4. Red nodes indicate up-regulation at protein level in RRMM compared to NDMM. Conversely, blue nodes indicated down-regulated proteins. Inset: PC metabolism pathways, showing expression correlation and proteomics up-regulation suggest active PC degradation in RRMM.

## Discussion

This study confirmed the feasibility of conducting concurrent lipidomics and proteomics profiling of freshly isolated plasma cells from patients with MM. We observed more lipidomic and proteomic differences between RRMM and NDMM, than between high and low risk MM based on the current R-ISS staging system. As an initial cross-validation, the proteomics data from our small pilot cohort was compared to a larger transcriptomics dataset for RRMM versus NDMM cases. This comparison revealed limited overlap at the transcript/gene level, likely due to the lower proteomics depth compared to transcriptomics. However, significant correlation was observed in the differential pathways at the transcript and proteome level, indicating agreement of our pilot cohort data with the larger transcriptome data. Together, these results confirm the feasibility of concurrent lipidomics and proteomics analyses from a single aliquot of one million plasma cells prepared from freshly collected bone marrow.

From both targeted and untargeted lipidomics, we observed significantly lower level of PC in RRMM compared to NDMM, and in high risk compared to low risk patients. Decreased PC was previously observed in MM cells compared to normal plasma cells [8]. Recently, Steiner *et al*. reported significantly lower circulating plasma levels of several PCs, and elevated lyso-PCs in RRMM compared to NDMM [27]. Hydrolysis of PCs by phospholipases generate lyso-PCs and a free fatty acid which could be further processed to generate lipid second messengers such as arachidonic acid, prostaglandins and leukotrienes [28]. These bioactive lipids play multiple roles in promoting cancer development and metastasis [29]. Interestingly, our transcriptomics network analysis of the larger independent cohort revealed high correlation of PC, arachidonic acid, prostaglandin metabolic pathways among RRMM. Furthermore, although the proteomic coverage of lipid metabolic enzymes was overall very limited, we found phospholipase B-like 1 gene product PLBD1 to be elevated in RRMM. The major cellular phospholipases that participate in signal transduction are PLA, PLC and PLD [28]. PLBD1 was recent identified from neutrophils as a phospholipase which removes fatty acids from either sn-1 or sn-2 positions [30]. Coupled with observed high level of transcripts in the arachidonic pathway, it is tempting to suggest that elevated PLBD1 levels contributes to MM progression and relapse by increasing arachidonic acids levels. Future studies in larger cohorts should examine this pathway.

We acknowledge that the small patient numbers in our study limit the broader applicability of the work, but in our small dataset, plasma cells from patients with RRMM appear to have a different lipidomic and proteomic profile when we compare with samples from NDMM. This is potentially clinically relevant, as patients who have relapsed disease experience poorer outcomes, with shorter periods of disease control than patients receiving front-line therapy at first diagnosis. The altered lipidomic and proteomic profile observed may reflect the clonal evolution that occurs in the malignant cells over time following serial chemotherapeutic challenges. To this end, it is interesting to note that PC is an important lipid in maintaining endoplasmic reticulum (ER) function, and that ER stress response pathways is implicated in the development of resistance to proteasome inhibitors in MM [31]. Further studies, with larger groups of patients will be beneficial in establishing the relationship between clonal evolution, subsequent lipidomic and proteomic changes. These results may enable personalized therapy selection, thereby improving patient outcomes.

In summary, we report the feasible concurrent lipidomic and proteomic analyses of purified plasma cells collected from a small cohort of multiple myeloma patients. As the goal was to determine the methodological feasibility and develop a suitable workflow, interpretation of the biological data from this study is limited by the small cohort size and possible confounders which were not considered. Nonetheless, in alignment with previous reports of reduced levels of PCs in MM (compared with healthy plasma cells), we observed reduced levels of several PCs in high risk MM and in RRMM. Furthermore, independent transcriptome data from a larger cohort corroborates altered PC metabolism in RRMM, and further suggest altered arachidonic acid and eicosanoid metabolism. We believe these preliminary observations warrants further exploration in a larger cohort, as these approaches are likely to provide valuable clinical insights into disease biology, as well as perhaps offer novel biomarkers for the prediction of disease kinetics.

## List of abbreviations

AA: arachidonic acid
FA: fatty acid
logFC: log fold change
MM: multiple myeloma
MTBE: methyl tert-butyl ether
NDMM: newly diagnosed multiple myeloma
PC: phosphatidylcholine
PE: phosphatidylethanolamines
R-ISS: Revised International Staging System
RRMM: relapsed/refractory multiple myeloma
SM: sphingomyelin

## Declarations

### Ethics approval and consent to participate

This study was approved by the PAH Human Research Ethics Committee (HREC/15/QPAH/442). Tissue banking was performed under the auspices of the Australasian Leukaemia and Lymphoma Group (ALLG) Tissue Bank.

## Acknowledgements

We would like to acknowledge the support of the Princess Alexandra Hospital Cancer Collaborative Biobank, and the clinical hematology laboratory staff who made this project possible, particularly Marlene Self, Donna Manning, Donna Cross and Sarah-Jane Halliday, as well as the QIMR Berghofer Medical Research Institute Proteomics Core Facility.

This project was funded, in part, by an Australian Cancer Research Foundation Grant “Diamantina Individualized Oncology Care Centre”. Lipidomics method development and analyses were enabled by a Translational Research Institute Spore Grant and Australian Research Council Discovery Project (DP160100224) and Future Fellowship (FT120100251) to MMH. KAM and JC are Queensland Health Junior Research Fellows.

## Authors’ contributions

KAM, MMH designed experiments. RJB, PMa, PMo facilitated access to suitable patients. JC, KAM recruited patients. HJ, JM performed experiments. AM conducted computational analyses. TS, FT, MRK contributed methodology. AM, JC, KAM, MMH drafted the manuscript. All authors approved the manuscript.

